# Histotripsy-initiated immune response synergizes with chemotherapy in a neuroblastoma murine model

**DOI:** 10.64898/2026.01.30.702878

**Authors:** Natalia Antonides-Jensen, Muskan Singh, Yuqing Xue, Fernando Flores-Guzman, Lydia L. Wu, Samantha S. Yee, Jacky Gomez-Villa, Timothy L. Hall, Mark A. Applebaum, Kenneth B. Bader, Sonia L. Hernandez

**Author notes:** These authors contributed equally to this work (Muskan Singh, Yuqing Xue). These authors share senior authorship (Kenneth B. Bader, Sonia L. Hernandez). **Corresponding author:** Kenneth Bader, University of Chicago, Department of Radiology 3835 S Cottage Grove Ave, Chicago, IL 60637.

## Abstract

High-risk neuroblastoma (NB) is a pediatric malignancy associated with metastases and an immunosuppressive tumor microenvironment. Standard-of-care treatments like chemotherapy are often ineffective, which motivates the investigation of adjuvant approaches. Histotripsy is a noninvasive focused ultrasound therapy that ablates tissue through the mechanical action of bubble clouds. In addition to disruption of the targeted tumor, non-targeted lesions exhibit growth delay after the histotripsy procedure. The primary hypothesis of this study was histotripsy-induced shifts in the tumor microenvironment will improve the response of metastatic NB to chemotherapy. Female A/J mice flanks were inoculated bilaterally with 1×10⁶ Neuro-2a cells. Histotripsy was applied to one tumor (200-500 mm³), with or without concurrent administration of liposomal doxorubicin (LDOX). The contralateral tumor served as a model of non-targeted distal metastases. Following treatment, tumors were monitored indefinitely for growth, or assessed after 5-7 days with flow cytometry, single-cell RNA sequencing, and immunohistochemistry. Histotripsy alone delayed the growth of treated and contralateral tumors relative to controls (*p* = 0.01 and *p* < 0.0001, respectively) and increased CD8⁺ T and CD11b^+^ cells (*p* < 0.05 for both comparisons). Further, NB cells in targeted and contralateral tumors exhibited a decrease in *Myc* expression and cell-cycle activity, and upregulation of interferon and apoptosis pathways. Histotripsy combined with LDOX had the longest delay in tumor growth (*p* < 0.01) and greatest expression of CD8⁺ and MOMA staining. These findings indicate that histotripsy induces a systemic antitumor immune response that potentiates chemotherapy efficacy in this model of metastatic NB.

**Significance:** Mechanical ablation with histotripsy drives systemic antitumor immunity, reshapes the tumor microenvironment, and enhances chemotherapy efficacy in a syngeneic model of metastatic, high-risk neuroblastoma.

## Introduction

Neuroblastoma (NB) is the most common extracranial solid tumor in children. Though the tumor cells are of neural crest origin, neuroblastoma generally arises in the adrenal gland.^1^ Of particular concern is high-risk NB, which has a five-year, event-free survival rate of 50%^2^ despite an aggressive treatment regimen that includes chemotherapy, surgery, autologous stem cell transplant, and immunotherapy.^1,2^ The poor response may be attributed in part to the extensive spread of the disease, as 70% of high-risk patients have metastases at the time of diagnosis.^3^ Further, the NB microenvironment remains hostile to endogenous immune elements via M2-polarized macrophages and myeloid-derived suppressor cells that interfere with therapeutic strategies.^4^ An approach that reprograms the microenvironment of primary and metastatic lesions and increases the efficacy of therapeutics is therefore needed for high-risk NB.

A potential method to target the tumor microenvironment is histotripsy, a focused ultrasound therapy that ablates tissue mechanically.^5^ Ablation is achieved via the application of pressure pulses with sufficient tension (> 25 MPa peak negative pressure) to generate bubble clouds spontaneously but controllably within the focal zone.^5^ The repeated mechanical stress of bubble cloud oscillations reduces the targeted region to acellular debris.^6^ Histotripsy has been cleared by the U.S. FDA to treat liver lesions in the adult population, and trials are underway to establish its safety and efficacy for targets in the kidney (NCT05820087) and pancreas (NCT06282809). There are several features of histotripsy that make it attractive for pediatric applications, including its non-invasive nature, lack of ionizing radiation, and lack of increased potential for metastatic disease.^7^ Further, histotripsy has been shown to induce a potent abscopal effect in other tumor models,^8,9^ and from a case report for a patient with multifocal lesions in the liver.^8–10^ Abscopal effects are hypothesized to originate from adaptive and innate immune responses. Specifically, histotripsy was shown to generate an increased concentration of CD8+ T cells relative to other ablation methods in a murine model of hepatocellular carcinoma.^11^ Shifts in natural killer cells following histotripsy exposure have also been observed, a primary immune target for NB.^12,13^

The capacity of histotripsy to generate an abscopal effect may sensitize the resistant NB microenvironment to conventional treatment approaches, such as chemotherapy. This premise is consistent with a previous study that found histotripsy induces two fundamental changes in high-risk NB: 1) increased expression of tumor necrosis factor alpha (TNF-alpha), and 2) an acute vascular dilation that lead to the mitigation of hypoxia.^12^ The former finding suggests histotripsy induces innate immunity through the action of natural killer cells.^14^ Adaptive responses could not be evaluated in those studies due to the immunocompromised nature of the model. The observed shifts in vascularity, however, suggested that the delivery of therapeutic agents may be enhanced via histotripsy. Indeed, prior studies have demonstrated histotripsy enhances the efficacy of therapeutic drugs or thrombo-occlusive disease.^13^ These effects may be accentuated for chemotherapy via the mitigation of hypoxia, a marker associated with a reduction in cytotoxic cell death.^15,16^

In this study, the effect of histotripsy on targeted and distal tumors was investigated with a syngeneic NB model. The primary hypothesis was that histotripsy would alter the immune cell composition in targeted and distal tumors, inducing an abscopal effect and delaying tumor growth.

## Materials and Methods

### Experimental Overview

This study consisted of two experimental components: (1) Evaluation of local and systemic immunological effects of histotripsy, and (2) Assessment of the therapeutic interaction between histotripsy and liposomal doxorubicin (LDOX), a chemotherapy agent investigated in pre-clinical neuroblastoma studies.^16^

For the immune response experiments, randomization within blocks was used to assign tumors to one of three groups: 1) untreated, 2) histotripsy, or 3) contralateral (the untreated tumor within a treated mouse). Within each group, a subgroup of mice was monitored for tumor growth until terminal endpoint criteria were met. For the other subgroup, animals were sacrificed 5-7 days post-treatment and tumor samples analyzed with flow cytometry, immunohistochemistry, single-cell RNA sequencing (scRNA-seq).

For the combination treatment experiments, randomization within blocks was used to assign animals to one of four groups: 1) control, 2) LDOX, 3) histotripsy, or 4) histotripsy combined with LDOX. Similarly, a subgroup of mice in each group was followed longitudinally to assess tumor growth and immunohistochemistry analysis. A separate subgroup was sacrificed acutely to evaluate intratumoral drug distribution.

### Cell Culture

Neuro-2a cells (ATCC, Manassas, VA; cat. no. CCL-131) were cultured in RPMI-1640 medium (Gibco, Waltham, MA; cat. no. 11875-093) supplemented with 10% heat-inactivated fetal bovine serum (Gibco, Waltham, MA; cat. no. 16000-044) and 1% penicillin–streptomycin (Gibco, Waltham, MA; cat. no. 15140-122). Cells were detached using 0.05% trypsin-EDTA (Gibco, Waltham, MA; cat. no. 25300-054). Cultures were maintained at 37°C in a humidified atmosphere containing 5% CO₂ and routinely tested for mycoplasma contamination (Invivogen, San Diego, CA; cat. no. rep-mys-10).

### Syngeneic Model

All animal experiments were approved by the University of Chicago IACUC #72341. Approximately 1×10⁶ Neuro-2a cells were injected subcutaneously into each flank of 5 to 6-week-old female A/J mice (The Jackson Laboratory, Bar Harbor, ME; cat. no. 000646). Mice without tumor growth or with tumors that ulcerated (0.5 cm diameter) prior to treatment were excluded from the study. Tumor length (*l*), width (*w*), and height (*h*) were measured with calipers, and the corresponding tumor volume was calculated with the ellipsoidal formula: 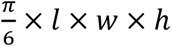. A maximal combined maximum tumor volume of 2000 mm^3^ was approved to follow the growth of both targeted and contralateral tumors after treatment. Animals were monitored routinely and removed immediately per IACUC guidelines. Treatment was administered when individual tumors reached 200-500 mm^3^, approximately 1-2 weeks after injections.

### Histotripsy

A diagram of the setup used to administer histotripsy is shown in Fig. 1. Histotripsy exposures were administered to target tumors in murine subjects under general anesthesia induced via ketamine. Prior to the treatment, the skin overlying the tumor was depilated using a commercial chemical depilatory agent (Nair, Church & Dwight, Ewing Township, NJ). Therapeutic ultrasound pulses were generated by a custom-designed 1.5-MHz focused transducer (20 mm focal distance, 45 mm outer diameter) driven by a custom-built Class D amplifier. Characterization of the acoustic field was performed using a fiber-optic probe hydrophone. Under linear acoustic propagation (−10 MPa), the −6 dB width for the peak pressure amplitude distribution for the measurement was 2.17 mm (acoustic axis), 0.73 mm (major axis), and 0.49 mm (minor axis).

**Figure 1.**
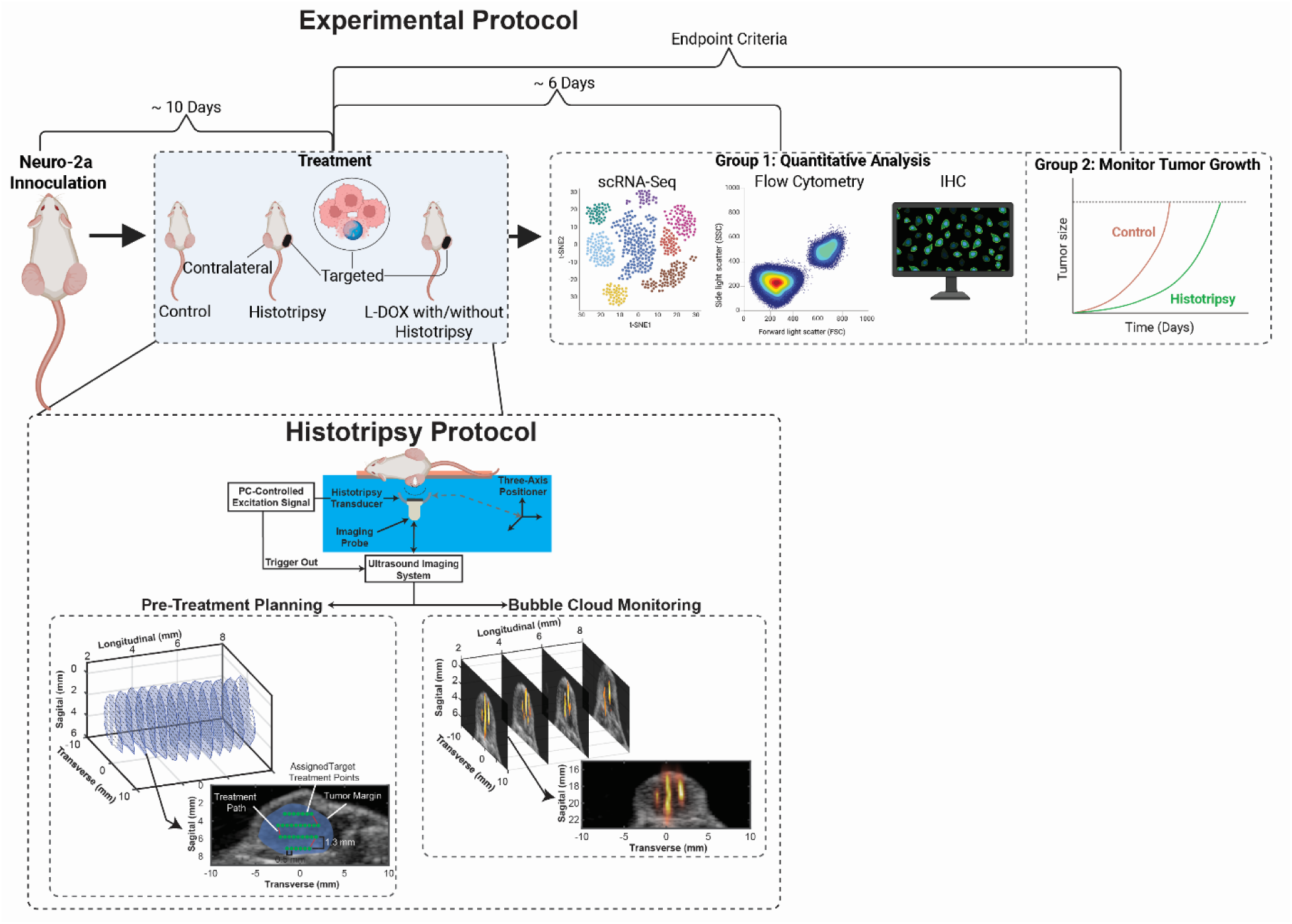
Schematic of the experimental workflow. Mice bearing bilateral Neuro-2a tumors received either no treatment, LDOX, the application of histotripsy to one tumor, or histotripsy and LDOX. To assess immune cell infiltration in treated and contralateral tumors, some mice were sacrificed 5-7 days post-treatment for immunohistochemistry, flow cytometry, and single-cell RNA sequencing analysis. Other mice were followed until terminal endpoint criteria were met. (Bottom) Ultrasound images acquired in 1-2 mm increments along the cranial-caudal axis of the tumor were used to reconstruct a volumetric treatment grid. A PC-controlled, three-axis motorized positioner was used to apply histotripsy pulses uniformly throughout the treatment volume. During treatment, bubble-cloud activity was monitored using real-time B-mode imaging. The same probe was used to collect acoustic emissions to form passive images and provide further confirmation of bubble cloud activity.

Anesthetized animals were positioned prone over a custom 3D-printed tray containing a centrally located circular cutout. This tray was placed within a tank filled with reverse osmosis–purified, 0.2 μm–filtered, degassed (pO₂ < 20%) water. The tumor was aligned within the cutout such that it was submerged during treatment. The water temperature was actively regulated and maintained at 37.0 ± 0.5 °C using a custom-built temperature controller (ITC-308, Inkbird, Shenzhen, China). An imaging probe (L11-5v, Verasonics, Kirkland, WA) aligned coaxially and confocally with the histotripsy transducer controlled by a research ultrasound system (Vantage 128, Verasonics, Kirkland, WA) was used to visualize the tumor. A custom MATLAB script (v2019b, MathWorks, Natick, MA) was used to acquire ultrasound images of the tumor at 1–2 mm intervals along its caudal–cranial axis. The images were interpolated to reconstruct a volumetric model of the tumor with a 0.5 mm interslice resolution. Within each reconstructed plane, a spatial treatment grid was defined with 0.5 mm lateral and 0.43 mm elevational resolution that covered 80% of the total tumor volume. These grid points defined the path through which the histotripsy focus was steered using motorized positioning stages (Velmex Inc., Bloomfield, NY) at a speed of 0.5 mm/s. Ultrasound pulses of ∼one cycle duration (0.67 µs) were applied at a rate of 100 Hz, corresponding to the application of 200 histotripsy pulses per millimeter of scanned tumor. To ensure consistent bubble activity throughout treatment and account for attenuation, an automated algorithm adjusted the electrical excitation based on the depth of penetration. Overall, the pulse peak negative pressure was 39.23 ± 4.68 MPa, and treatment duration 23.17 ± 12.93 min.

During insonation, bubble-cloud generated acoustic emissions were acquired passively with the L11-5v imaging probe and processed with a pth-root algorithm to form passive images (Fig. 1).^17,18^ Acoustic emissions serve as a surrogate for the mechanical strength of bubble cloud activity.^19,20^ The images were analyzed qualitatively to confirm bubble cloud activity (and therefore the potential for mechanical ablation) was applied consistently throughout the histotripsy exposure.

### Chemotherapy administration

LDOX (FormuMax Scientific, Sunnyvale, CA; cat. no. F10101-NC-5) was administered intravenously via the tail vein at 1 mg/kg using insulin syringes under brief restraint. The formulation was diluted with sterile PBS on the day of treatment and adjusted to a total injection volume of 100 µL per mouse. For mice in the combination group, LDOX was administered 1-4 hours before histotripsy treatment. Note the dose of LDOX (1 mg/kg) was selected to result in minimal disruption of tumor growth relative to untreated controls but still provide a detectable fluorescent signal.^21^

### Tumor growth curves and AUC analysis

For mice enrolled in survival studies, tumor size was measured daily until the time of sacrifice. Tumor growth curves were generated in GraphPad Prism (v10.6.1; GraphPad Software, Boston, MA). Tumor growth was plotted for each tumor until the tumor reached 300% of its day 0 volume or until the mouse met ethical tumor burden criteria (any single tumor exceeding 1000 mm³ or total tumor burden exceeding 1500 mm³). The area under the curve (AUC) for tumor volume over time was calculated using the trapezoidal rule.

### Tumor harvest for single cell suspension

Upon sacrifice, tumors were excised and the skin tissue was dissected away. Scissors were used to mince the tumor into chunks before mixing with 3 to 4 mL of PBS within a Falcon tube on ice strained through a 40 µm nylon mesh cell strainer (Thermo Fisher Scientific, Waltham, MA; cat. no. 08-771-1). The tumor chunks were collected and placed into a Falcon tube with freshly prepared 4 mL of RPMI with 5% FBS with 1 mg/mL collagenase A (Sigma-Aldrich, St. Louis, MO; cat. no. C9891) and 100 µL of 1 mg/mL DNase I (Roche, Basel, Switzerland; cat. no. 10104159001). This mixture was incubated for 30 minutes at 37°C. The Falcon tube was gently inverted every 5 minutes. After incubation, the solution was filtered through a 40 µm nylon mesh cell strainer and centrifuged for 4 minutes at 1000 rpm. The pellet was resuspended in 2.5 mL of PBS and 2.5 mL ACK Lysing Buffer was added (Quality Biological, Gaithersburg, MD; cat. no. 118-156-101), incubated at room temperature for 30 seconds to remove red blood cells. After centrifugation, the supernatant was removed, and the pellet was resuspended in PBS. Trypan Blue was used to check cell viability and cell count before proceeding with Dead Cell Removal Kit (Miltenyi Biotec, Bergisch Gladbach, Germany; cat. no. 130-090-101) per manufacturer’s instructions. Cell preparations with under 70% cell viability were excluded from analysis. The average viability percentage did not differ significantly amongst groups via (state test here): untreated 81±7.5% (N = 5), histotripsy 89 ± 5% (N = 2), and contralateral 74± 6% (N = 2).

### scRNA-sequencing

Library preparation and sequencing were performed by the University of Chicago Genomics Core (RRID:SCR_019196). Chromium Next GEM Single Cell 3′ Reagent Kits v3.1 (10x Genomics, Pleasanton, CA) were used for library generation and NovaSeq X-10B-100 (Illumina, San Diego, CA) for sequencing.

### Data processing and quality control

Barcoded libraries sequenced with 80 bp paired-end reads (barcode: 12 bp cell barcode, 8 bp UMI) and 60 bp (3′ end of transcript) served as the input for data processing. The Cell Ranger package (v8.1.0; 10x Genomics, Pleasanton, CA) was used to process the scRNA-seq FASTQ files. The “cellranger count” function, with default command settings, aligned FASTQ files to the mouse GRCm39 (refdata-cellranger-arc-GRCm39-2024-A) reference genome and generated an expression matrix with gene read counts per cell. The filtered count matrices were read into the R package Seurat (v5.3.0; Satija Lab, New York, NY). Cells with less than 200 or more than 7,500 expressed genes, or more than 10% mitochondrial content were removed. Samples that had abnormal cell density distributions (more than 30% low-quality cells) were discarded. The data were normalized using the “NormalizeData” function with the “LogNormalize” method. To reduce background signals before integration, “FindVariableFeatures” was used to identify the top 2,000 feature genes with the highest variability. Samples were then integrated with the “IntegrateData” function using these feature genes to remove batch effects. The “ScaleData” function with default settings was applied to the integrated dataset, followed by Principal Component Analysis (PCA) using the “RunPCA” function.

### Clustering and annotation

To downscale and identify clusters, Unified Flow Approximation and Projection (UMAP)^22^ was performed with the “RunUMAP” function in Seurat, using an optimized number of principle components (PCs) defined by Elbow Method. Marker genes for each cluster were identified using the “FindAllMarker” function with logfc.threshold = 0.25 and min.pct = 0.1 as the threshold.^23^ The cell type of each cluster was manually annotated by comparing marker genes with published canonical cell-type signatures.^24^ Predictions from SingleR (v 2.6.0)^25^ were used as supplementary guidance. Bar plots of cell proportions were made in GraphPad Prism (v10). Cell cycle scores were calculated, and cell cycle phases were assigned to tumor cells using the “CellCycleScoring” function in Seurat, with gene lists for G2/M and S phases converted from human to mouse orthologs prior to use.

### Differential gene expression and gene set enrichment analysis

Differentially expressed genes between groups were identified using the “FindMarkers” function in Seurat with the Wilcox-Limma test method. Here, the required gene expression was 5% of cells in each group. To identify enriched pathways, genes with adjusted *p* value < 0.05 and average log2 fold change > 0.25 or < −0.25 were pre-ranked based on log2 fold change. Gene set enrichment analysis (GSEA) was performed using clusterProfiler (v4.12.6; He lab, Guangzhou, China).^26^ Gene ontology (GO) Biological Process enrichment was performed using “gseGO” function, and hallmark pathways enrichment was conducted using “GSEA” function with MSigDB Hallmarker gene set for mouse.

### Flow cytometry

A portion of the single-cell tumor suspension was processed with flow cytometry. Cell suspensions were fixed and permeabilized with CytoFast Fix/Perm Buffer Set (BioLegend, San Diego, CA; cat. no. 426803) and stained with the panel of antibodies described in Sup. Table 1. Analyses were performed on a BD LSRFortessa with High Throughput Sampler (BD Biosciences, Franklin Lakes, NJ) and analyzed using FlowJo software (v10.9.0; BD Life Sciences, Ashland, OR).

An overview of gating strategies is shown in Sup. Fig. 3. The following gating strategy was performed to visualize CD4 T cells: The whole cell population was first gated on singlets, second on debris exclusion, third on CD3 positive cells, fourth on double positive cells for CD4 APC and CD3 PE, and fifth on IFN gamma FITC with CD4 APC double positive cells. The following gating strategy was performed to visualize CD8⁺ T cells: The whole cell population was first gated on singlets, second on debris exclusion, third on CD3 positive cells, fourth on double positive cells for CD8 APC and CD3 PE, and fifth on IFN gamma FITC with CD8 APC double positive cells. On an independent sample, a sixth gate was applied for Perforin FITC and CD8 APC. The following gating strategy was performed to visualize B cells, NK cells, macrophages, and CD11b^+^ cells: The whole cell population was first gated on singlets, the second gate on debris exclusion and enrichment of white blood cell populations, and the third gate on the population of interest: B cells gated on CD19 PE positive cells, NK cells gated on CD56 PE positive cells, and macrophages on F4/80 eFluor570 positive cells. Individual samples were stained with antibodies labeled with fluorescent dyes PE and FITC that required a compensation matrix. Final counts were processed in GraphPad Prism.

### LDOX extraction and quantification in tumor tissue

Assessment of intratumoral LDOX was performed as previously described.^21^ Tumor tissues were kept on dry ice throughout the experiment. Approximately 0.1-0.15 grams of frozen tumor tissue were weighed and transferred into individual Lysing Matrix D tubes (MP Biomedicals, Santa Ana, CA; cat. no. 6913100). For each 0.1 g of tissue, 500 µL of nuclear lysis buffer (0.25 M sucrose, 5 mM Tris-HCl, 1 mM MgSO₄, 1 mM CaCl₂, pH 7.6) was added. Samples were then processed using FastPrep-24 5G (MP Biomedicals, Santa Ana, CA; cat. no. 116005500) with the following settings: 6 m/sec for 40 seconds. After homogenization, tubes were centrifuged at 3,000 rpm for 4 minutes at 4°C. The supernatant was collected, and 200 µL of each sample was aliquoted. For 200 µL volume of tumor extract, 100 µL of 10% Triton X-100, 200 µL of water, and 1 mL of acidified isopropanol (0.75 N HCl) were added. Tubes were briefly vortexed, then stored overnight at −20°C. The following day, samples were brought to room temperature, vortexed for 45 seconds, and centrifuged at 2,000 g for 15 minutes. The resulting supernatant was collected and stored at −80°C until analysis.

For analysis, samples were brought to room temperature, and a volume of 100 µL from each sample was plated in triplicate into an opaque 96-well plate. Fluorescent signal was measured at 470 nm excitation and 590 nm emission using a SpectraMax i3x plate reader (Molecular Devices, San Jose, CA; cat. no. 10014-924). A standard curve for LDOX quantification was prepared by adding 2 µL of LDOX stock solution (4 mg/mL) to 400 µL of solvent, resulting in a final concentration of 20,000 ng/mL. This extract was serially diluted in acidified isopropanol using a 1:3 dilution scheme. Concentrations ranging from 246.91 ng/mL to 0.000465 ng/mL were selected from these dilutions to generate the standard curve to ensure all tumor readings were within the standard curve. To account for autofluorescence, the mean signal of untreated control samples was subtracted for samples that included LDOX. The corresponding baseline adjusted signals were then correlated with the standard curve to determine the concentration of drug in Graphpad Prism. The calculated amount of LDOX was normalized to the mass of tumor loaded.

### Assessment of LDOX distribution

A subgroup of animals was sacrificed 24 hours after treatment, and a ∼7 mm x ∼5 mm section was resected from each tumor along the sagittal plane and mounted without being fixed or stained. Once mounted, slides were stored at −70°C until imaging. Slides were imaged using a Nikon Eclipse Ti2 fluorescent microscope (Nikon Instruments, Tokyo, Japan) and then returned to cold storage, spending no more than 1 hour at room temperature prior to image collection. All images were acquired as dual-channel composites (brightfield and 550 nm fluorescence) at 20X magnification. Five images were collected for each slide at random locations of tissue. Microscope gain, exposure, and laser intensity settings were kept constant throughout all image collection.

Collected images were analyzed in ImageJ (National Institutes of Health, Bethesda, MD) to quantify the percentage of area covered by fluorescence. Composite files were separated into brightfield and fluorescent channels. The brightfield channel was used to identify and mask tissue voids within the corresponding fluorescent image. Fluorescent images were then filtered using the “Threshold” function in ImageJ to reduce auto fluorescent signal. The threshold value was determined by averaging the luminance level at which untreated samples exhibited >0.01% area coverage. Following filtering, the percent area coverage of fluorescence for each slide was quantified using the “Measure” function in ImageJ.

### Immunohistochemistry

For paraffin-embedded slides, tissue sections underwent deparaffinization and rehydration, followed by heat-mediated antigen retrieval. For CD8 staining, antigen retrieval was performed using a 1-hour incubation low pH IHC Antigen Retrieval Solution (Invitrogen, Waltham, MA; cat. no. 00-4955-58). For MOMA staining, antigen retrieval was carried out using EDTA-Tris buffer for 1 hour. Slides were then cooled to room temperature and incubated in 0.5% Triton X-100 for 10 minutes. After washing, blocking was performed with CAS-Block (Thermo Fisher Scientific, Waltham, MA; cat. no. 008120) for 1 hour. Primary antibodies were diluted in CAS-Block and incubated overnight at 4°C: anti-CD8 alpha (1:100, Abcam #ab217344) and anti-MOMA-2 (1:400, Abcam #ab33451). The sections were subsequently incubated with horseradish peroxidase-conjugated secondary antibodies for 1 hour: Rabbit anti-Rat IgG Alexa Fluor 488 (1:250, Invitrogen #A21210) and Chicken anti-Rabbit IgG Alexa Fluor 488 (1:250, Invitrogen #A21441). A final wash was followed by nuclear staining with DAPI (Vector Laboratories, Burlingame, CA; cat. no. H1800). Slides were imaged using an Eclipse Ti2 microscope (Nikon, Tokyo, Japan).

The same staining protocol was applied to fresh-frozen slides from LDOX and histotripsy combination experiments, including permeabilization, blocking, primary and secondary antibody incubation, and DAPI staining. Steps specific to paraffin-embedded tissue were not performed for fresh-frozen slides, including deparaffinization, rehydration, and heat-mediated antigen retrieval.

### Statistical analysis

All statistical analyses were performed using GraphPad Prism (v10.6.1, GraphPad Software, Boston, MA) or RStudio (v2024.12.1+563, Posit PBC, Boston, MA). Data distribution was assessed prior to analysis, and nonparametric methods were used when normality assumptions were not met. Tumor growth over time was summarized using AUC, calculated via the trapezoidal rule, and group differences were assessed using Welch’s one-way ANOVA followed by Games-Howell post hoc tests to account for unequal variances and sample sizes. Flow cytometry data were analyzed using one-way ANOVA with Tukey’s post hoc test. For LDOX extraction and intratumoral concentration measurements, group differences were assessed using the Kruskal-Wallis test followed by Dunn’s post hoc test for pairwise comparisons. Differences in LDOX spatial distribution were analyzed using a linear mixed-effects model with repeated measures, with Dunn’s test used for post hoc comparisons. For scRNA-seq analyses, differences in the proportion of immune cell subtypes between experimental groups were assessed using the Kruskal-Wallis test to evaluate overall differences across groups. Given the small sample size no pairwise comparisons were performed. Differences in Myc expression and cell cycle phase scores between tumor groups were assessed using two-sample t-tests with Benjamini-Hochberg correction for multiple testing.

## Results

### Administration of histotripsy

The therapy transducer generated sufficient pressure to initiate a bubble cloud for all cases as evidenced by detection via hyperechogenicity on standard B-mode imaging and strong acoustic emissions (Fig. 1 and Sup. Video 1). The presence of bubble clouds based on these imaging markers coincided tumor damage based on gross and histological analyses of targeted tumor (Sup. Fig. 1). No treatment-related issues or adverse responses were observed in the subjects during or after the procedure. Notably no increase in tumor ulceration frequency was observed for histotripsy-exposed tumors (ulcerated tumors were excluded prior to treatment to minimize variation).

### Histotripsy alone delays tumor growth and has an abscopal effect

Tumor volumes measured with calipers over time are shown in Fig. 1A. The tumor growth curves suggest delayed tumor growth in subjects exposed to histotripsy relative to untreated controls. This qualitative assessment was supported by AUC analysis, which was increased for both histotripsy-treated and contralateral tumors relative to untreated controls (*p* = 0.013 and *p* < 0.0001, respectively; Welch’s ANOVA with Games-Howell post hoc; Fig. 1B). No significant difference was observed between histotripsy-treated and contralateral tumors (*p* = 1.00). These findings provide evidence of an abscopal effect, as tumor growth was inhibited in non-targeted, contralateral tumors.

**Figure 2.**
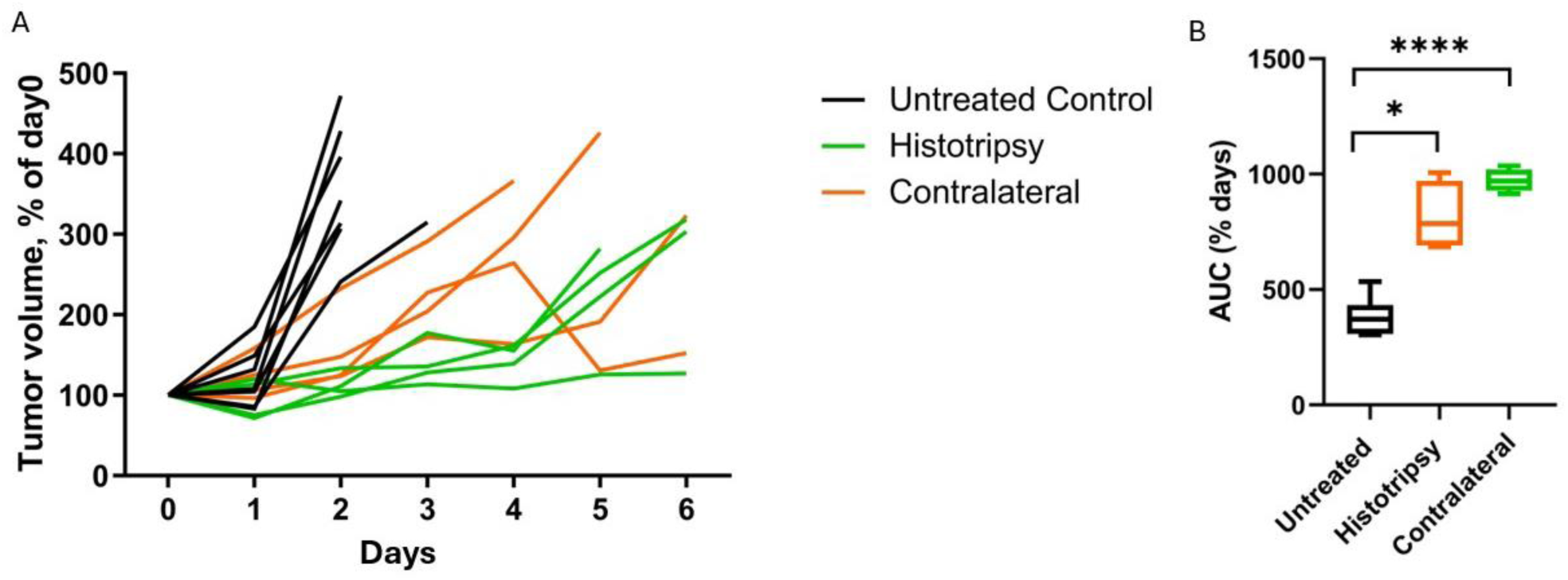
Histotripsy delays tumor growth and has an abscopal effect. **A,** Normalized tumor volume over time for untreated controls (black; n = 7), histotripsy (green; n = 4), and contralateral (orange; n = 4). Histotripsy was applied on day 0. Histotripsy (green; n = 4) slowed tumor progression relative to untreated controls (black; n = 7). **B,** Area under the curve (AUC) for each treatment arm. Welch’s ANOVA with Games–Howell post hoc testing indicated the AUC values were increased for both the histotripsy and contralateral groups relative to untreated controls (p = 0.013 and *p* < 0.0001, respectively). There were no differences in AUC values between histotripsy and contralateral tumors. *, *p* ≤ 0.05; ****, *p* ≤ 0.0001.

### Histotripsy alone elicits both innate and adaptive immune responses

Global shifts in the immune cell landscape are a potential mechanism for the reduction in tumor growth under histotripsy. To obtain a complete overview of cell populations in response to histotripsy, three untreated control tumors and two of histotripsy-treated and corresponding contralateral side of tumors were analyzed by scRNA-seq. In total, four cell types could be labeled including neuroblastoma (NB) tumor cells, T cells, myeloid cells, and fibroblasts (Sup. Fig. 2A). Cell populations were annotated through canonical gene markers (Sup. Fig. 2B). High resolution analysis was achieved via selection of cells within an immune subgroup that expressed the *Ptprc* gene (encoding mouse CD45 protein). These cells were then re-clustered, and immune cell subtypes were defined by their expression of hallmark genes (Sup. Fig. 2C).

The myeloid and T cell subtypes identified are indicated in Fig. 3A. Compared to the untreated tumors, treated and contralateral tumors showed changes in their immune cell populations (Fig. 3B). A Kruskal-Wallis test indicated differences between these groups for cytotoxic CD8^+^ T-Cells (*p =* 0.03; Fig. 3C), and anti-inflammatory, pro-tumor M2 Macrophages (*p =* 0.03; Fig. 3D, left). No differences were noted between groups for M1 Macrophages (*p =* 0.27; Fig. 3D, right).

**Figure 3.**
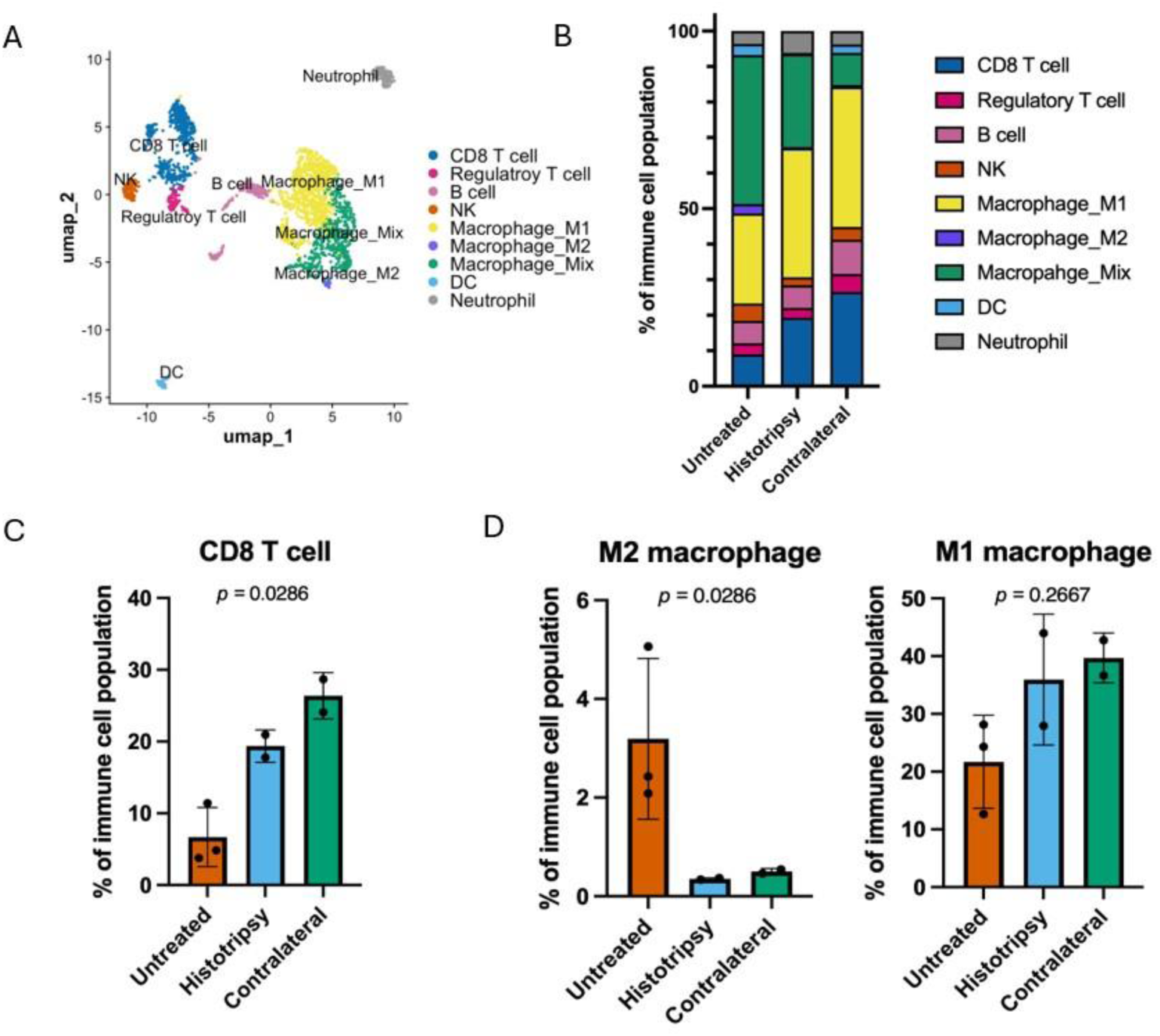
Histotripsy induces innate and adaptive immune activation in both targeted and contralateral tumors. **A,** Annotated UMAP of CD45^+^ immune cells. **B,** Cell-type proportions**. C,** Proportion of cytotoxic CD8⁺ T cells for each group. A difference was observed between all groups (*p =* 0.03; Kruskall-Wallace test). **D,** Myeloid cell composition between groups. A difference was observed between all groups for M2 macrophages (*p =* 0.03; Kruskall-Wallace test, left), but not M1 macrophages (*p =* 0.27; Kruskall-Wallace test, right).

These findings were further supported by flow cytometry. In tumors targeted by histotripsy, the fraction of CD3⁺ CD8⁺ T cells was increased for treated (3.77 ± 2.3%) and contralateral (4.05 ± 1.8%) groups relative to untreated controls (1.26 ± 0.99%; *p* < 0.05 for both comparisons; Welch’s ANOVA with Games-Howell post hoc; Sup. Fig. 3). No difference between histotripsy-treated and contralateral tumors was detected for CD3⁺ CD8⁺ T cells. Differences were also noted for CD11b⁺ cells in treated (24.03 ± 13.96%) and contralateral groups (27.24 ± 14.82%) relative to untreated controls (7.7 ± 3.7%; *p* < 0.05 for both comparisons; Welch’s ANOVA with Games-Howell post hoc; Sup. Fig. 3). No differences were noted between histotripsy-treated and contralateral tumors for CD11b⁺ cells. No differences were noted between experimental arms for other markers investigated with flow cytometry (Sup. Fig. 3). Notably, no differences in the fraction of natural killer cells were noted between arms for either flow cytometry or scRNA-seq. Together, these results indicate histotripsy upregulates innate and adaptive immune responses in this model of high-risk NB.

### Histotripsy alone increases non-viable tumor area and immune cell recruitment in both treated and contralateral tumors

Upon histological analysis, large regions of non-viable tumor tissue were present in histotripsy-treated tumors and their contralateral counterparts 6 days after treatment, but not in untreated controls (Fig. 4A, pink areas). Cells with immune-like morphology were observed along the borders of these non-viable regions (Fig. 4A, green). Immunohistochemistry indicated an infiltration of CD8⁺ T cells along the boundary between viable and non-viable tumor (Fig. 4B, arrow, boundary marked by line). In contrast, the macrophage marker MOMA staining was more prevalent in non-viable tumor regions (Fig. 4C, green, arrow, boundary marked by line).

**Figure 4.**
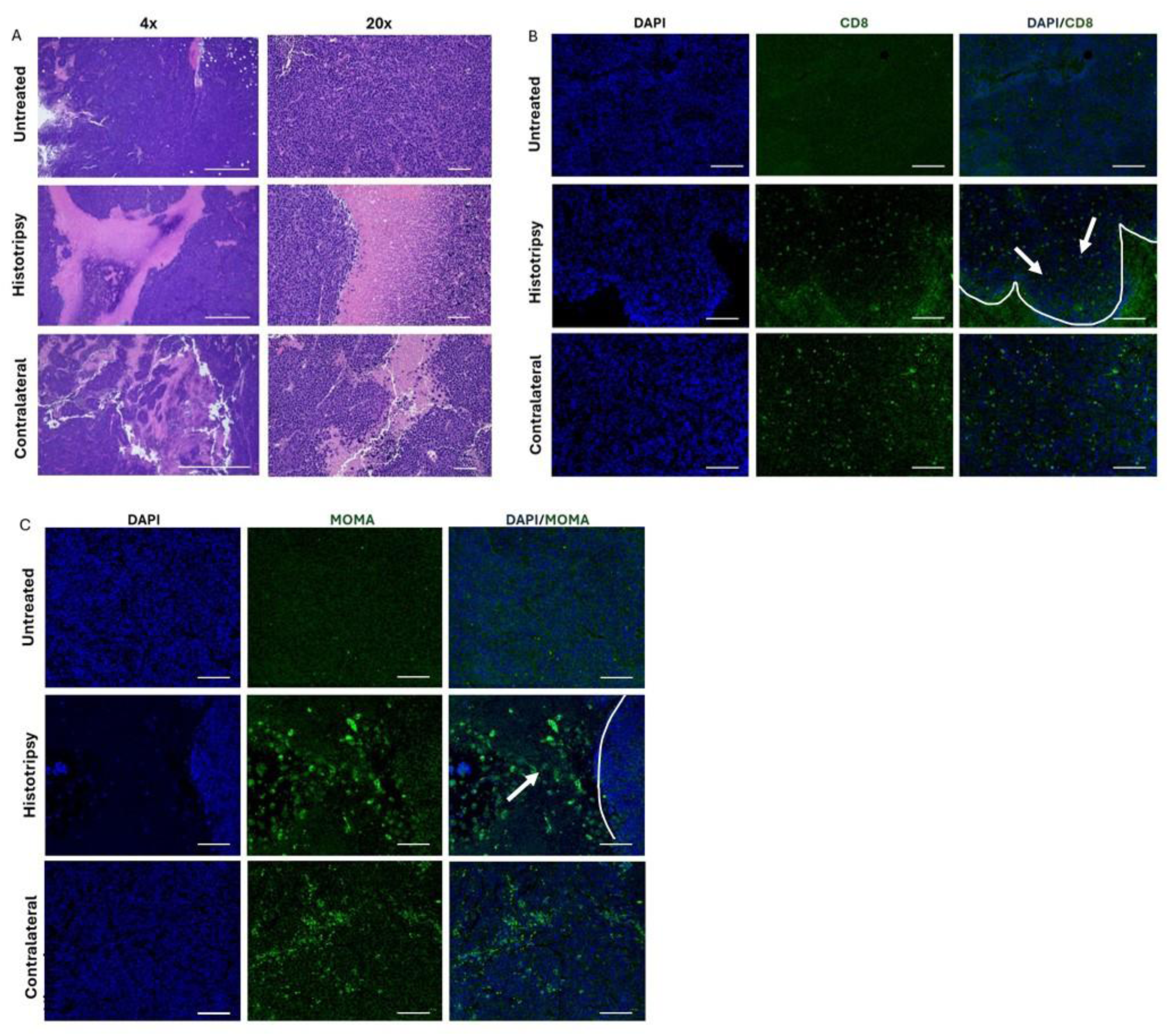
Histotripsy increases non-viable tumor area and immune cell recruitment in both targeted and contralateral tumors. **A,** Representative H&E images showing large regions of non-viable tumor tissue in histotripsy-treated tumors and in contralateral tumors, which were absent in untreated controls. Immune-like cells were observed along the borders of these non-viable regions. 4X and 20X magnification. Scale bars: 1000 µm and 100 µm (left to right). **B,** Representative CD8 immunohistochemistry demonstrating increased localization of CD8⁺ T cells at the interface between viable and non-viable tumor areas in both histotripsy-treated and contralateral tumors. 20X magnification. Scale bar: 200 μm. CD8⁺ T cells marked by arrow, boundary marked by line. **C,** Representative MOMA immunohistochemistry showing increased macrophage presence at the same interface in treated and contralateral tumors compared to untreated controls. 20X magnification. Scale bar: 200 μm. Macrophage cells marked by arrow, boundary marked by line

### Histotripsy induces transcriptional and cell cycle changes in neuroblastoma cells

In addition to changes in the immune landscape, histotripsy was also found to alter the NB cell cluster. The oncogene *Myc*, a key oncogene that drives proliferation in neuroblastoma, was found to have a reduced expression in histotripsy and contralateral tumors relative to untreated controls (*p <* 0.001 for both; t-test with Benjamini-Hochberg adjustment; Fig. 5A). Further, Myc expression was decreased in contralateral tumors relative to those exposed to histotripsy (Fig. 5A). Cell cycle phase is indicated in Fig. 5B. The overall G2/M score of treated and contralateral groups was decreased compared to untreated control, indicating lower cell proliferation activity in groups associated with histotripsy exposure (*p* < 2e-6*, p* = 5e-16, respectively; t-test with Benjamini-Hochberg adjustment; Fig. 5C). Further supporting these findings, MYC, G2M, and E2F MsigDB hallmarks were all identified as decreased in treated and contralateral compared to control tumors (Fig. 5D). While expression of Myc and cell cycle was decreased in treated and contralateral tumors, we found that these tumors had a significant increase in MSigDB hallmarks of interferon gamma, interferon alpha, and apoptosis pathways, consistent with the increase in immune activation identified in the immune cell cluster (Fig. 5E). We also compared the gene set enrichment using gene ontology (GO) database and found similar enrichment of pathways of immune activation (Fig. 5E).

**Figure 5.**
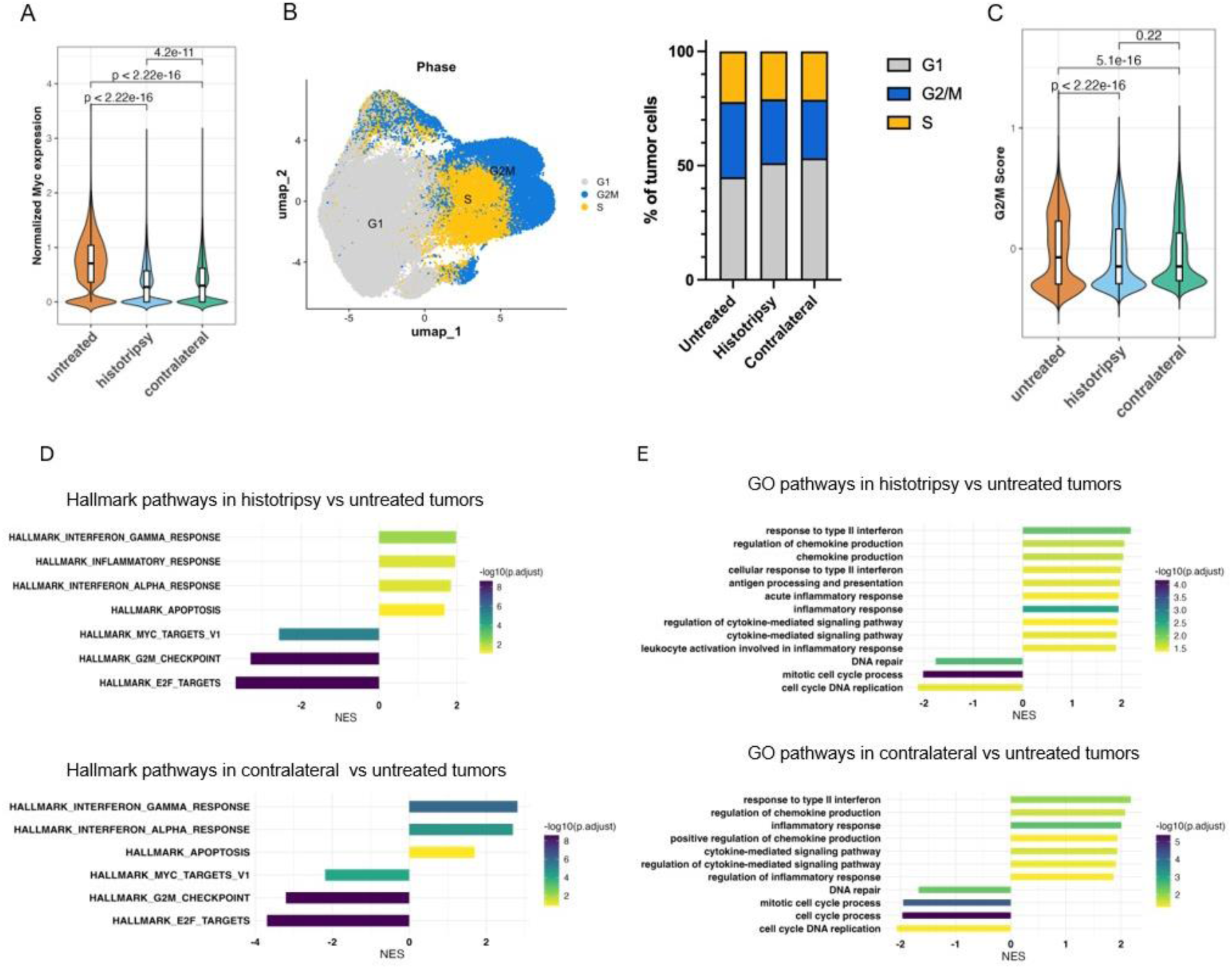
Histotripsy reduces tumor cell proliferation locally and distally. **A,** *Myc* activity in Neuro-2a tumor cell clusters, showing significantly decreased *Myc* pathway scores in both histotripsy-treated (n = 3) and contralateral tumors (n= 2) compared with untreated controls (n = 2; *p <* 0.001 for both; t-test with Benjamini-Hochberg adjustment). **B,** Cell-cycle phase assignment generated to illustrate the distribution of G1, S, and G2/M phase cells across untreated, treated, and contralateral tumors. **C,** Quantification of G2/M cell-cycle scores demonstrating reduced proliferative activity in histotripsy-treated and contralateral tumors relative to untreated controls t (*p <* 0.001 for both; t-test with Benjamini-Hochberg adjustment). **D,** Differential pathway activity in tumor cells, with treated and contralateral tumors showing decreased *Myc* and cell cycle expression and increased MSigDB hallmarks of interferon gamma response, interferon alpha response, and apoptosis (NES, normalized enrichment score). **E,** Gene Ontology enrichment analysis demonstrating upregulation of pathways associated with immune activation in treated and contralateral tumors.

### Histotripsy synergizes with liposomal doxorubicin (LDOX)

The tumor volume normalized to day 0 (i.e., day of treatment) and area under the curve (AUC) is shown in Fig. 6A for arms with and without LDOX, and with and without histotripsy. Tumor growth appeared most stunted for tumors targeted with histotripsy combined with LDOX (n = 6) relative to untreated controls (n = 7), LDOX alone (n = 8), or histotripsy alone (n = 4). Analysis based on AUC indicated that overall cumulative tumor burden was reduced for histotripsy combined with LDOX relative to all other groups (p < 0.0001 vs. untreated controls; p < 0.001 vs. LDOX alone; p < 0.001 vs. histotripsy alone; Welch’s ANOVA with Games–Howell post hoc). For contralateral tumors, the combination treatment delayed tumor growth compared with untreated controls and LDOX alone (Fig. 6B; both p < 0.01). Although a trend toward delayed tumor growth was observed for the combination treatment relative to histotripsy alone (AUC = 1441.5 ± 314.2 % days vs. 971.8 ± 49.5 % days, respectively), this difference did not reach statistical significance (p = 0.09).

**Figure 6.**
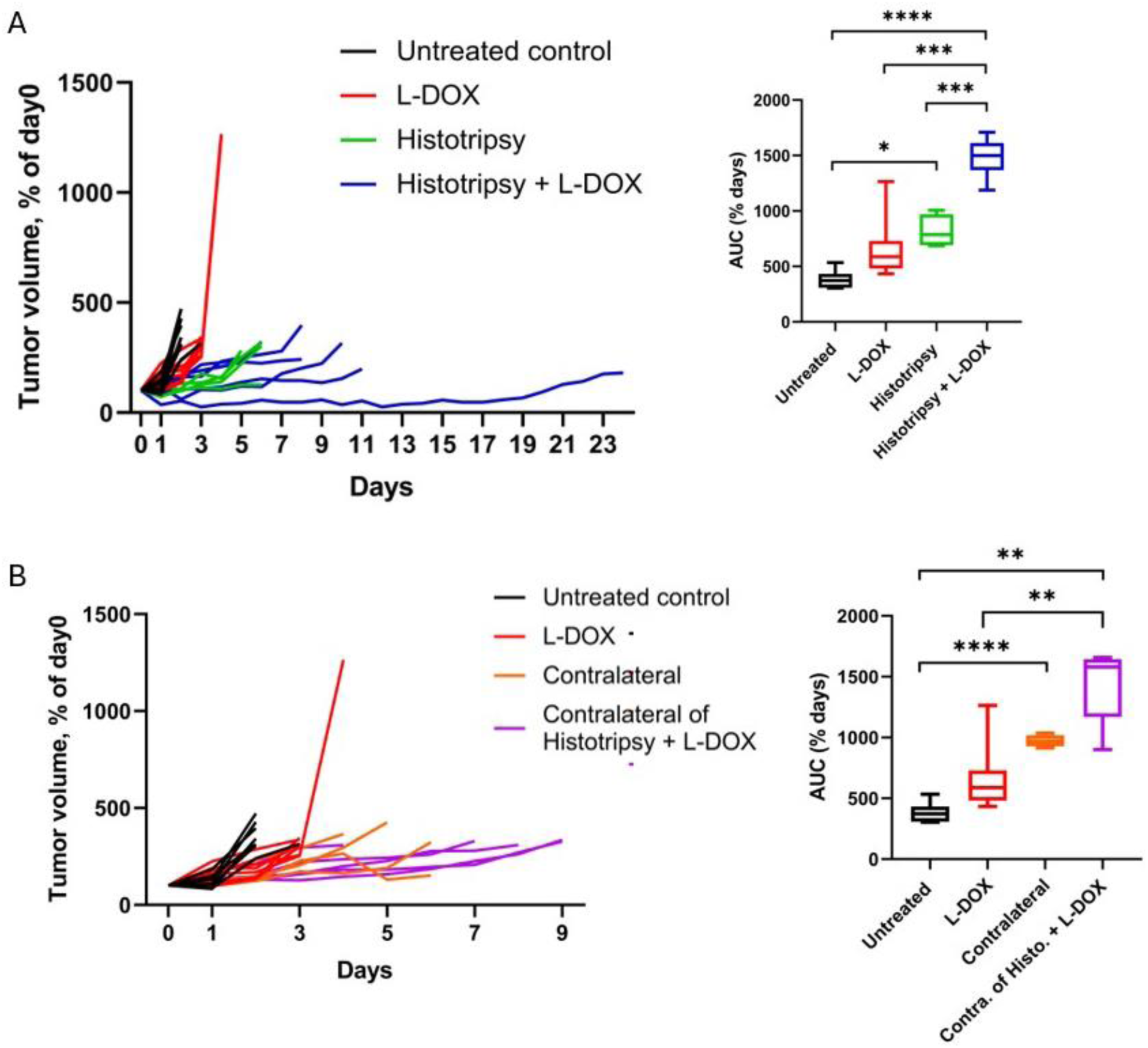
Histotripsy combined with LDOX mitigates tumor growth. **A,** Normalized tumor growth curves for histotripsy combined with LDOX (blue; n = 6), untreated controls (black; n = 7), LDOX (red; n = 8), and histotripsy (green; n = 4). The AUCs were reduced when histotripsy was combined with LDOX compared with untreated controls (p < 0.0001), LDOX alone (p < 0.001), or histotripsy alone (p < 0.001; Welch’s ANOVA with Games–Howell post hoc). **B,** Normalized growth curves for contralateral tumors for the combination treatment group (purple; n = 5), untreated controls (black; n = 7), LDOX (red; n = 8), and histotripsy (orange; n = 4). Combination treatment significantly delayed tumor growth compared with untreated controls and LDOX alone (both p < 0.01; Fig. 6B). Although a trend toward delayed tumor growth was observed relative to histotripsy alone, this difference was not statistically significant (p = 0.09). *, p ≤ 0.05; **, p ≤ 0.01; ***, p ≤ 0.001; ****, p ≤ 0.0001.

### Histotripsy does not increase short-term uptake of LDOX

Outcomes for the extraction of LDOX from tumors 24 hours after histotripsy exposure are indicated in Fig. 7A and overall tumor coverage in Fig. 7B. Similar findings were observed for the area coverage of LDOX. Although some individual cases showed strong LDOX coverage after histotripsy in both targeted (3 of 5) and contralateral tumors (2 of 3), this pattern was not significant overall (Fig. 7C; p > 0.05). Similar findings were observed when the total tumor coverage by histotripsy was decreased to 40% tumor coverage (Sup. Fig. 5).

**Figure 7.**
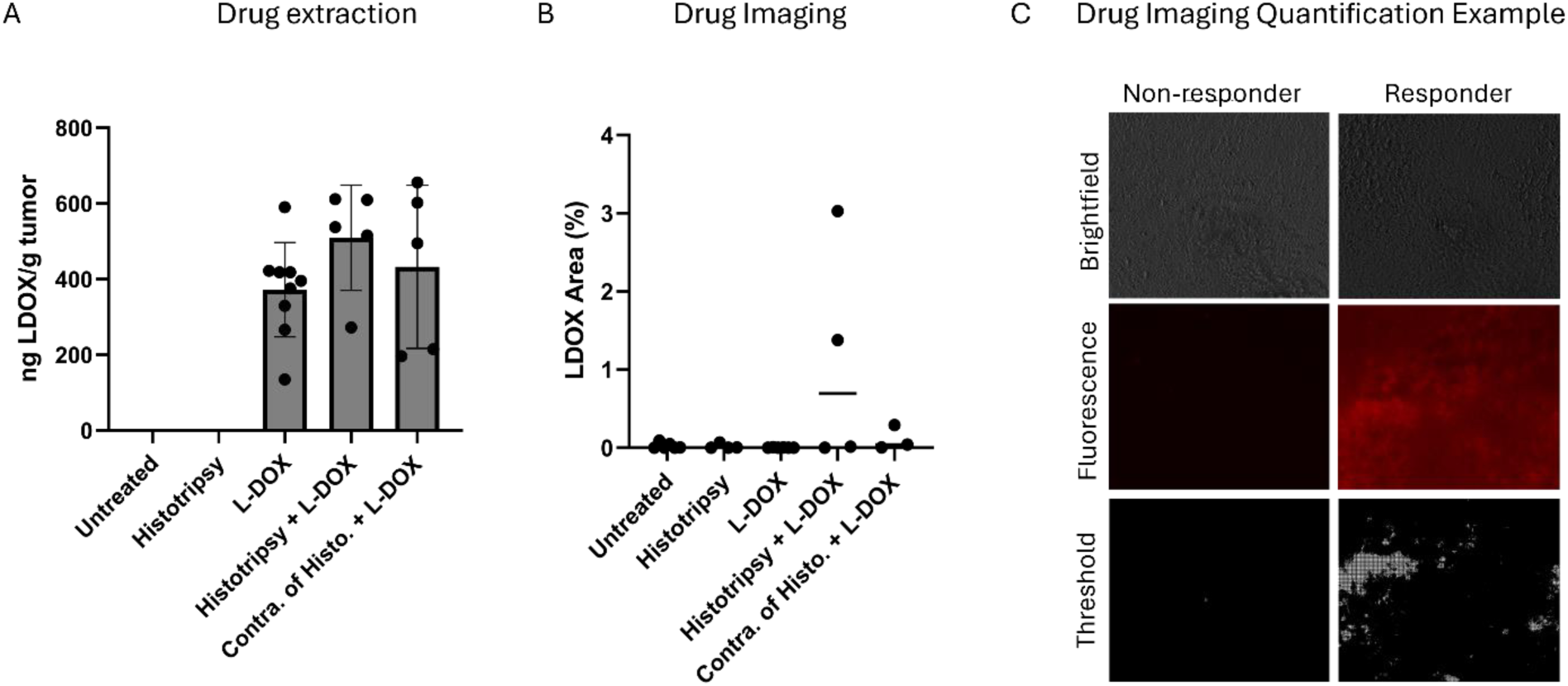
Histotripsy does not increase short-term uptake or distribution of LDOX in tumors. **A,** Quantification of LDOX extraction from tumors 24 hours after treatment (untreated, *n* = 8; histotripsy, *n* = 3; LDOX, *n* = 9; histotripsy + LDOX, *n* = 5; contralateral of histotripsy + LDOX, *n* = 5). No differences were observed among groups that included LDOX (*p* > 0.05; Kruskal–Wallis test with Dunn’s post hoc test). **B,** Fractional tumor coverage by LDOX (untreated, *n* = 6; histotripsy, *n* = 4; LDOX, *n* = 4; histotripsy + LDOX, *n* = 3; contralateral of histotripsy + LDOX, *n* = 5). No differences were observed among groups that included LDOX (*p* > 0.05; Kruskal–Wallis test with Dunn’s post hoc test). **C,** Representative brightfield and fluorescence images from the histotripsy + LDOX group. The combination treatment showed cases with no detectable LDOXsignal (non-responders, *n* = 2 of 5) and substantial LDOX signal (responders, *n* = 3 of 5).

### Combined histotripsy and LDOX treatment increase immune cell infiltration in both treated and contralateral tumors

Given the pronounced synergistic effect observed despite minimal changes in drug uptake or distribution, additional analysis of immune cell recruitment was performed. Immunohistochemistry staining indicated histotripsy and LDOX treated tumors express similar levels of expression for CD8 and MOMA (Fig. 8A and 8B). These markers were concentrated in regions bordering viable and non-viable tumor. The prevalence of CD8 and MOMA was accentuated for tumors treated with histotripsy combined with LDOX, as well as their contralateral counterparts.

**Figure 8.**
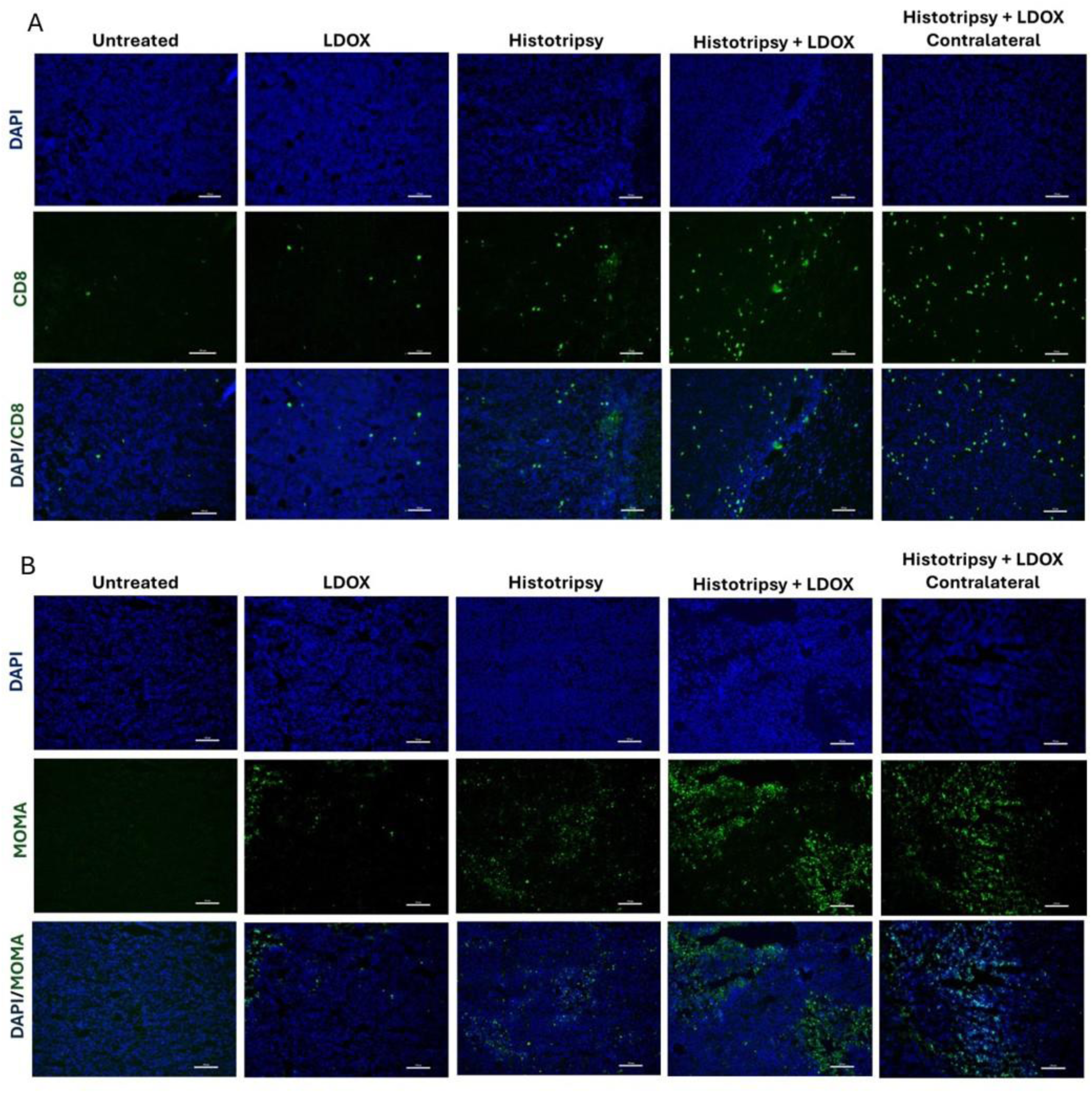
Histotripsy combined with LDOX increases immune cell infiltration in treated and contralateral tumors. **A,** Representative images of CD8 immunohistochemistry showing modest increases in CD8⁺ T cells at the interface between viable and non-viable tumor regions in tumors treated with histotripsy alone or LDOX alone compared to untreated controls. Tumors treated with the combination of histotripsy and LDOX exhibited further increases in CD8⁺ T cells, which were also observed in contralateral tumors. **B,** Representative images of MOMA immunohistochemistry demonstrating similar patterns for macrophages, with modest increases in tumors treated with histotripsy or LDOX alone and more pronounced infiltration in combination-treated tumors and their contralateral counterparts. All images: 20X magnification; Scale bar: 200 μm.

## Discussion

Current treatment paradigms for high-risk NB are inhibited by its immunologically cold nature and chemoresistivity.^27,28^ This study evaluated whether histotripsy ablation could be counter these features in multifocal NB via single-tumor ablation. Histotripsy exposure alone resulted in a reduction in tumor growth for treated and contralateral tumors compared to untreated controls (Fig. 2), consistent with an abscopal effect. This observation may be attributed in part to two factors: changes in the immune landscape and metabolism of NB cells. Targeted and contralateral tumors showed higher proportions of cytotoxic CD8⁺ T cells and myeloid cells based on scRNA-seq (Fig. 3), flow cytometry (Sup. Fig. 3) and immunohistochemistry (Fig. 4). Further, differences in protumor M2 macrophages were noted between groups, with a trend for reduction in targeted or contralateral tumors (Fig. 3D). Similar histotripsy-induced abscopal effects have been noted in pre-clinically in multiple cell lines, and select patients with multi-focal hepatocellular carcinoma.^29,30^ The cell line used in this study (Neuro-2a) forms tumors with low baseline PD-L1 expression and limited T-cell infiltration, consistent with the immunologically “cold” nature of high-risk NB.^31,32^ Therefore, these data may represent a worse-case scenario for the abscopal effect, through further data is needed for confirmation. It should be noted no differences were noted in natural killer cell activity between groups, a primary immune target for NB.^12^ Tumors in this study were analyzed 5-7 days after histotripsy exposure, whereas a prior investigation noted natural killer cell activity may be most active at earlier timepoints.^33^ Differences in the tumors models may contribute to this discrepancy, but motivate future studies to assess morphological changes in the immune landscape

Beyond increases in immune cell infiltration, a significant finding for this study was the impact of histotripsy on NB cell metabolism. Both treated and contralateral tumors exhibited downregulation of Myc and cell-cycle pathways, along with upregulation of interferon and apoptosis pathways (Fig. 5). Together, these data indicate a global reduction in the proliferative activity of the NB cells. It remains unclear whether these changes in the metabolism of NB cell were driven by increased immune cell infiltration or other factors. A prior study indicated an upregulation of TNA-alpha in histotripsy treated tumors, which can sensitize cells to apoptotic stimuli associated with punitive changes in metabolic activity.^34,35^ Nevertheless, tumor cell metabolism and immune infiltration are tightly interconnected, and the application of histotripsy is associated with initiation of this bidirectional feedback loop.

These effects were further enhanced when histotripsy was combined with LDOX. In targeted tumors, the combination therapy had a reduced tumor growth relative to all other treatment arms. Although contralateral tumors did not exhibit a statistically significant growth delay with combination treatment, a similar directional trend was observed. The lack of statistical significance in contralateral tumors may reflect the limited sample size. There are multiple factors that motivate a combination treatment. While histotripsy alone was able to drive an abscopal effect (Fig. 2), prior studies indicate the application of therapeutic ultrasound alone does not provide sustained tumor control for metastatic disease.^29^ Additionally, there are practical limitations to the application of a local therapy like histotripsy in metastatic diseases such as high-risk NB, including the potential for post-ablation syndrome and tumor targetability.^36,37^ Previous studies have shown that histotripsy can alter tissue in ways that enhance drug delivery, including vascular dilation and reduction of tumor hypoxia, a feature associated with decreased chemotherapy efficacy.^34,38,39^ A primary hypothesis of this study was therefore that histotripsy-induced changes in the tumor microenvironment would improve intratumoral drug delivery and kinetics. Interestingly, no acute increase in drug uptake or distribution was observed for histotripsy independent of the degree of expose (Fig. 7 and Sup. Fig. 5). Moreover, hallmark hypoxia pathways were upregulated in both treated and contralateral tumors compared to untreated controls (Sup. Fig. 4), contrary to previous findings showing reduced histologic evidence of hypoxia 24 hours after treatment.^34^ These differences may reflect the specific timepoint analyzed, the neuroblastoma tumor model used, or the parameters of the histotripsy source applied in this study. This discrepancy highlights the need for further investigation into the temporal and mechanistic effects of histotripsy on drug delivery and tumor biology.

Despite the lack of increased LDOX retention, the combination treatment provided the most effective tumor control (Fig. 6). The half-life for LDOX is just over a day, and tumor control for the combination treatment extended tumor control on average by 6.62 days for targeted tumors and 2.03 days for contralateral tumors relative to histotripsy alone.^40^ These findings suggest mechanisms other than the immediate additive effects of the two therapies. The greatest CD8⁺ T cell and macrophage infiltration occurred in the combination treatment group and extended to both treated and distant tumors (Fig. 8). The precise reason for the observed synergy is unknown. Apoptotic pathways initiated by LDOX and histotripsy may occur through orthogonal or complementary mechanisms. The generation of reactive oxygen species is a major intrinsic apoptotic pathway for doxorubicin, which can be generated during histotripsy.^41^ Caspase-3 is another intrinsic pathway for doxorubicin cell death and has been increased via histotripsy exposure in another NB model.^34^ Regardless of the precise mechanism, these findings suggest immune activation as a central mechanism driving the therapeutic synergy.

Several limitations of this study should be considered. All experiments were performed in a single subcutaneous Neuro-2a tumor model and all female mice, which does not fully recapitulate the biology or metastatic patterns of high-risk NB including as the involvement of bone marrow metastasis. A narrow range of histotripsy parameters was evaluated. There is increasing evidence that overtreatment with histotripsy (i.e., the number of pulses applied per unit volume) may mitigate the corresponding immunological effects.^33^ Studies with a histotripsy dose response are needed to outline treatment regimens that provide effective therapeutic delivery and response. Tumor genomic analysis with scRNA-seq was performed at a single time point (day 6). While earlier or later time points could provide additional insight into the temporal evolution of immune activation, this time point allowed both adaptive and innate immune responses to be captured. Finally, a fixed and low dose of LDOX was tested.^21^ While liposomal doxorubicin has been tested in clinical and pre-clinical studies, standard doxorubicin combined with multiple other forms of chemotherapy is more commonly used as part of the treatment platform.^16,42^ Together, these factors limit the generalizability of our findings and highlight the need for future studies across additional tumor models, metastatic sites, chemotherapy drugs, and histotripsy treatment conditions.

Nevertheless, these results suggest that histotripsy induces a robust global response driven by immune activation across both treated and distant tumors, which is further amplified in combination with LDOX. These findings support studying histotripsy in combination with immunotherapy, specifically due to histotripsy-induced immune activation, to further enhance systemic anti-tumor immunity in a NB model.

## Data Availability

The data generated in this study are available upon request from the corresponding author.

## Acknowledgements

This project is support in part by the Focused Ultrasound Foundation, Arms Wide Open Childhood Cancer Foundation, American Cancer Society (Grant # RSG-21-171-01-ET), and National Institutes of Health (Grant # R01 EB0352309). We acknowledge support provided by the UChicago Cytometry and Antibody Technology Core Facility (Facility RRID: SCR_017760) and Functional Geonomics Core Facility (RRID:SCR_019196).

---

**Supplementary Figure 1.**
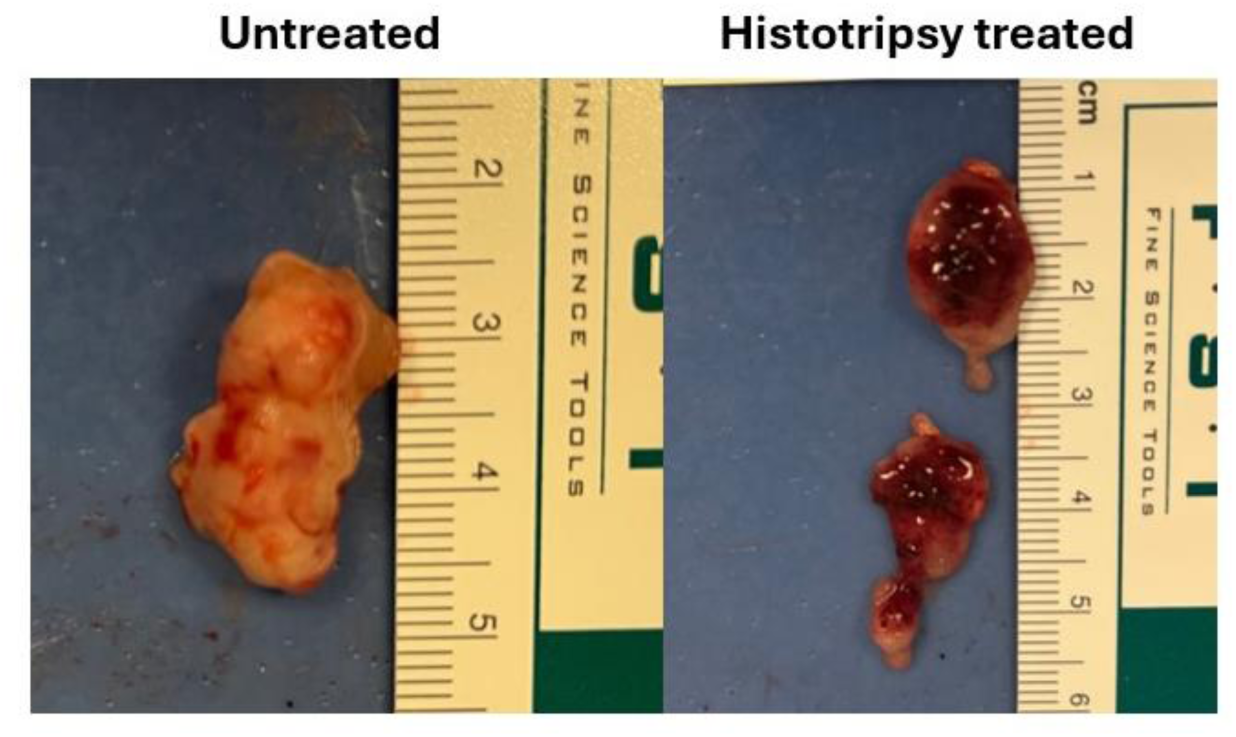
Representative images of untreated tumors and tumors treated with histotripsy (80% of tumor targeted). Treated tumor has been sectioned in half.

**Supplementary Figure 2.**
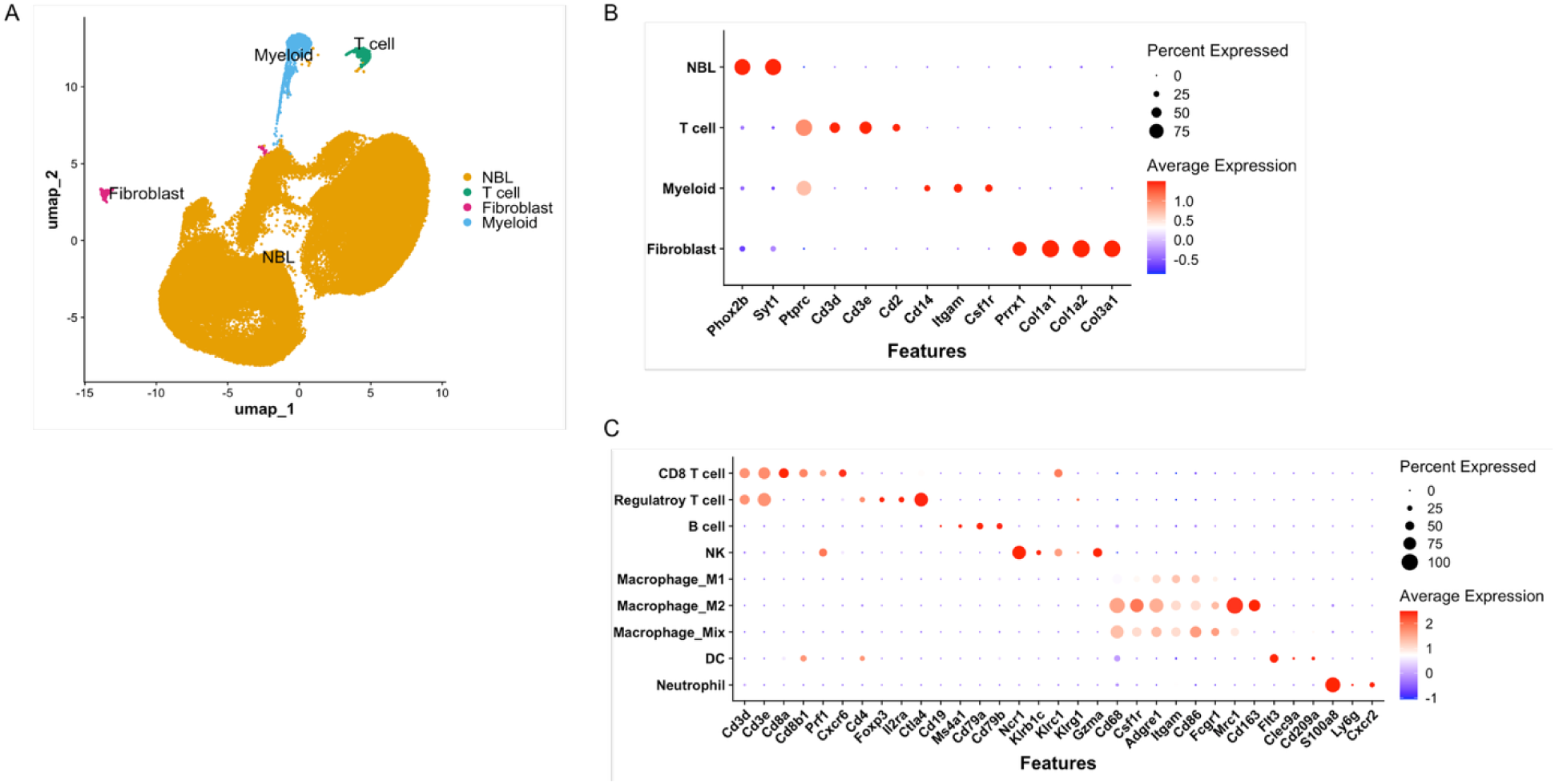
**A,** Annotated integrated uniform manifold approximation and projection (UMAP) of scRNA-seq from untreated (n = 3), histotripsy-treated (n = 2), and contralateral tumors (n = 2). Four major cell types were identified: neuroblastoma tumor cells (NBL), T cells, myeloid cells, and fibroblasts. **B,** Dotplot of key marker gene expression used for annotation in A. **C,** Dotplot of key marker gene expression used for annotation in Fig. 3A.

**Supplementary Figure 3.**
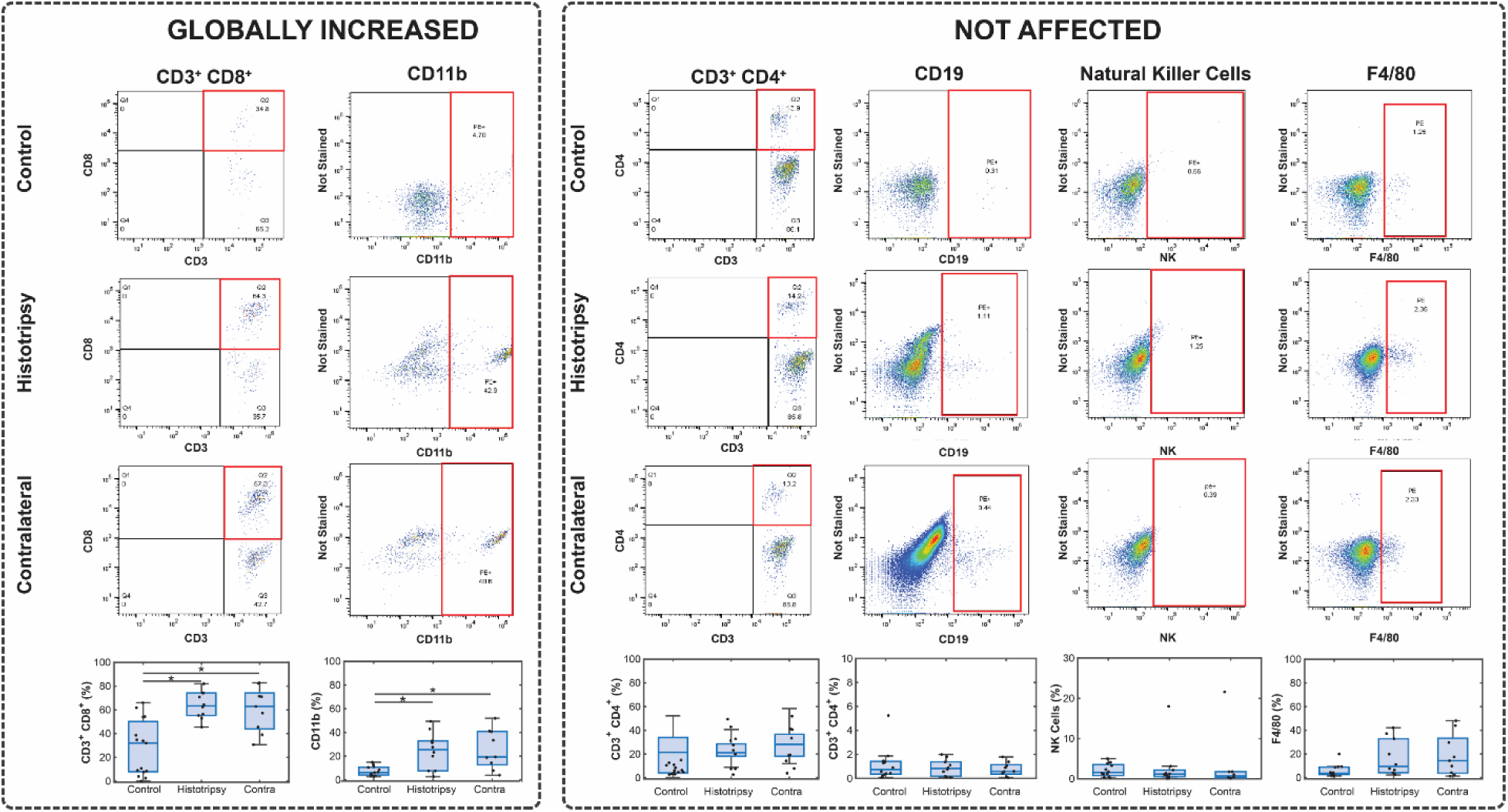
Flow cytometry analysis indicated histotripsy elicited both innate and adaptive immune responses. The percentage of CD3⁺CD8⁺ T cells was increased in histotripsy-treated and contralateral tumors compared with untreated controls (*p* < 0.05 for both comparisons, Welch’s ANOVA with Games-Howell post hoc). Similarly, CD11b⁺ cells were increased in histotripsy-treated and contralateral tumors relative to untreated controls (*p* < 0.05 for both comparisons, Welch’s ANOVA with Games-Howell post hoc). No significant differences were observed between histotripsy-treated and contralateral tumors for either CD3⁺CD8⁺ T cells or CD11b⁺ cells. No significant differences were observed across experimental groups for the other markers assessed with flow cytometry.

**Supplementary Figure 4.**
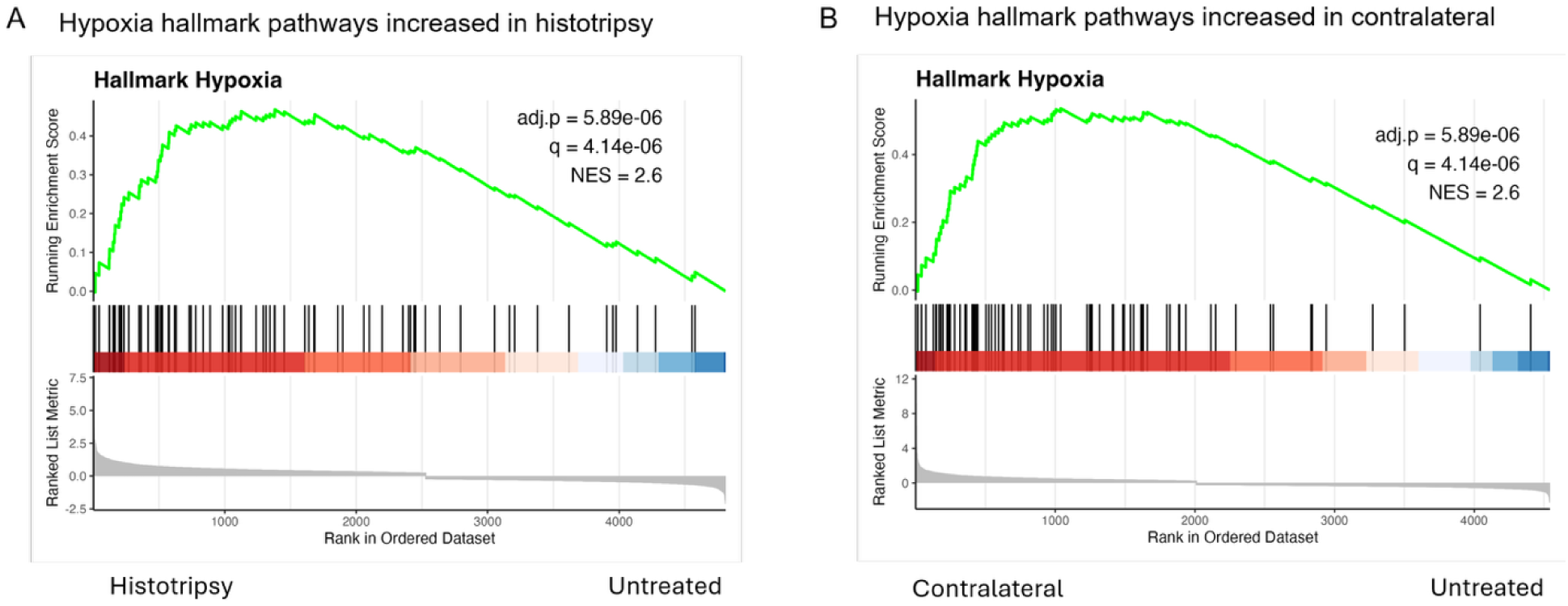
**A,** GSEA enrichment plot for hypoxia hallmark pathway for histotripsy-treated tumors compared with control. **B,** GSEA enrichment plot for hypoxia hallmark pathway in contralateral tumors compared with control.

**Supplementary Figure 5.**
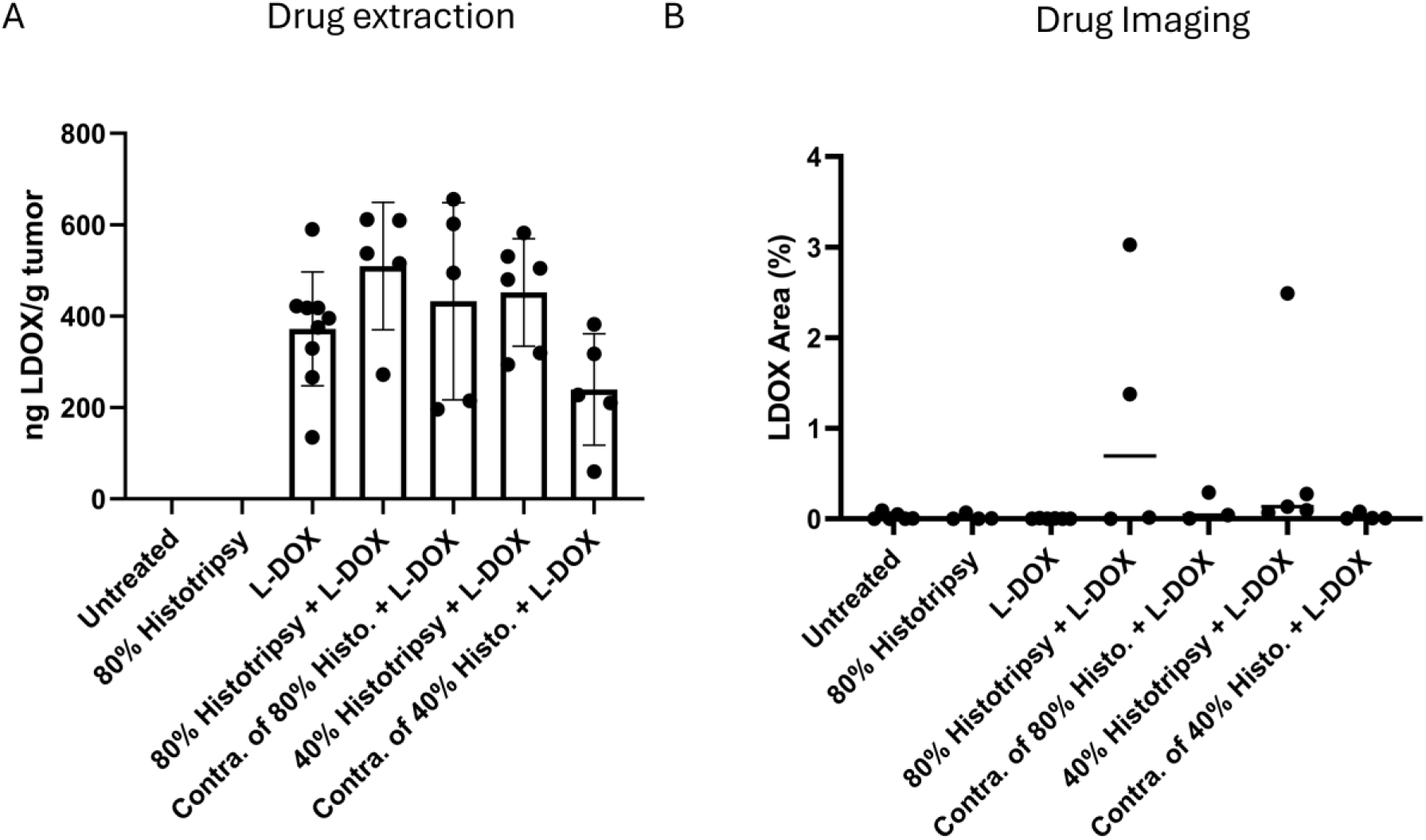
Histotripsy does not increase short-term uptake or distribution of L-DOX in tumors for 40% to 80% tumor coverage. **A**, Quantification of L-DOX extracted from tumors 24 hours after treatment (untreated, n = 8; 80% histotripsy, n = 3; L-DOX, n = 9; 80% histotripsy + L-DOX, n = 5; contralateral of 80% histotripsy + L-DOX, n = 5; 40% histotripsy + L-DOX, n = 6; contralateral of 40% histotripsy + L-DOX, n = 5). Tumors receiving combined histotripsy + L-DOX, regardless of total tumor coverage by histotripsy, showed no significant increase in L-DOX levels compared to L-DOX alone (unpaired t-test, *p* > 0.05). Contralateral tumors similarly showed no significant difference relative to L-DOX alone (unpaired t-test, p > 0.05). **B**, Assessment of the area of L-DOX distribution within tumors (untreated, n = 6; histotripsy, n = 4; L-DOX, n = 4; 80% histotripsy + L-DOX, n = 3; contralateral of 80% histotripsy + L-DOX, n = 5; 40% histotripsy + L-DOX, n = 5; contralateral of 40% histotripsy + L-DOX, n = 4). While individual histotripsy-treated tumors showed strong L-DOX coverage, there were no overall differences between groups (one-way ANOVA, p > 0.05).

**Supplementary Table 1.**
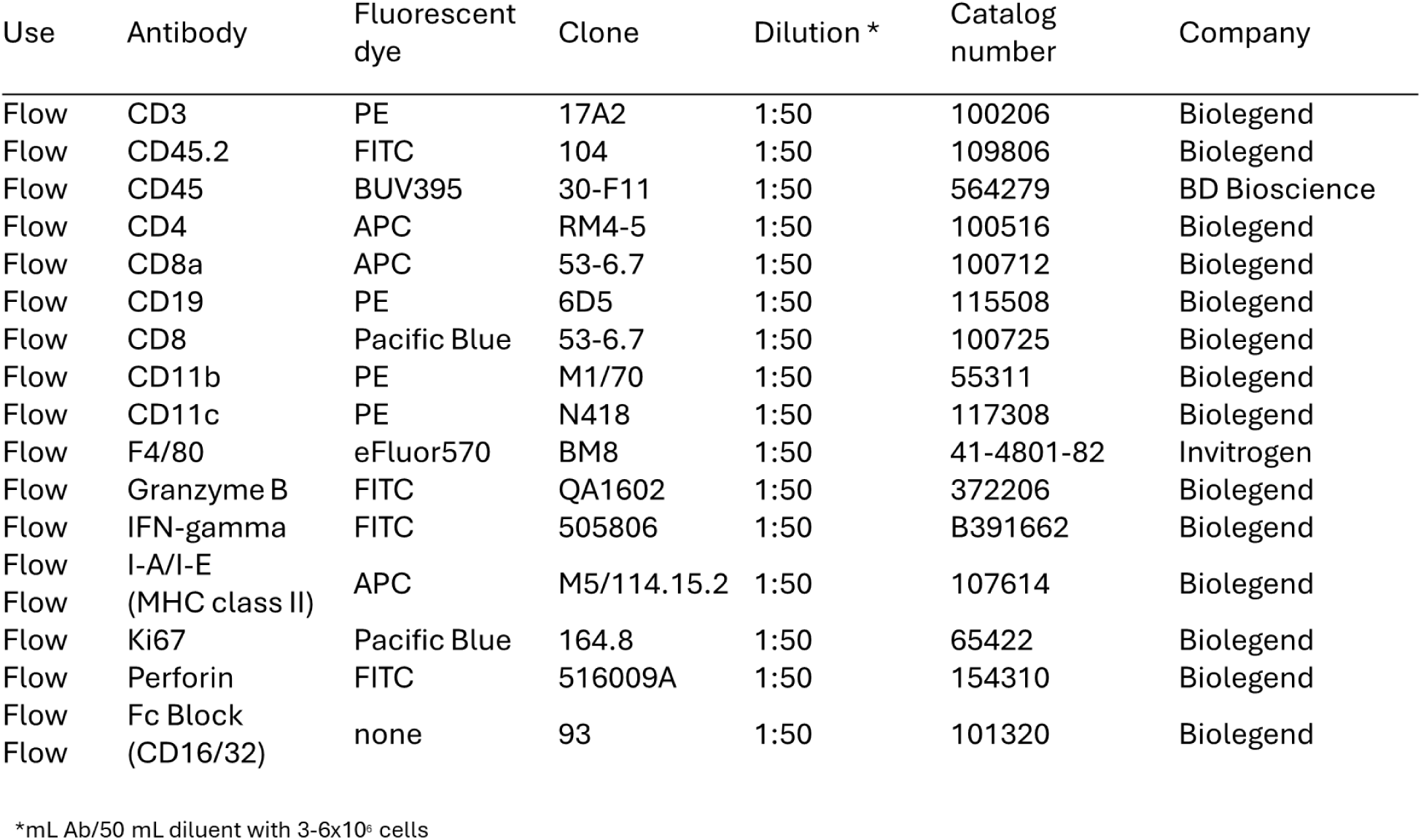
Panel of antibodies used to stain single-cell tumor suspensions for flow cytometry.

